# Methyl gallate Attenuates Post-Stroke Emotional and Cognitive Symptoms by Promoting Hippocampal Neurogenesis via PI3K/GSK3 and AMPK Signaling

**DOI:** 10.1101/2025.07.08.663507

**Authors:** Limin Xu, Feifei Chen, Haimei Fan, Longxing Zhang, Fei Yang, Jianyu Pang, Baolong Pan, Xuemei Wu

## Abstract

Restoring hippocampal neurogenesis is an effective strategy for post-stroke recovery. Methyl gallate (MET) exhibits neuroprotective properties. However, the effect of MET in improving brain functional recovery in the post-stroke depression (PSD) model and its underlying mechanism remains unknown. Single-cell data analysis showed that the cell types and molecular characteristics of PSD are similar to those of primary depression but exhibit weaker synaptic plasticity and stronger inflammatory signals. In addition, molecular docking studies revealed that MET exhibits a significant binding capacity with AMPK/GSK3, suggesting that MET mediates the neuroprotective effects of both. In this study, we created a post-stroke depression (PSD) model by performing physical restraint after ischemia and tested the treatment effects of MET. We observed that MET significantly attenuated PSD-induced depressive-/anxiety-behaviors associated with a reduction of stress hormone corticosterone and ACTH levels. Morris water maze and recognition task results indicate that MET can also alleviate cognitive impairments in the PSD model. In the hippocampus of the PSD model, MET improved the proliferation and differentiation of neural stem/progenitor cells (NSPCs). MET treatment significantly enhanced the activity of AMPK and decreased the activity of GSK3β. Furthermore, in primary neural progenitors under hypoxia, both the PI3K inhibitor LY294002 and the AMPK inhibitor compound C blocked the effects of MET to promote neural development. Animal experiments also confirmed that LY294002/compound C treatment could reduce the effects of MET in antidepressant behaviours. Taken together, our results indicate that PI3K, as well as AMPK-mediated adult neurogenesis, is restored by MET to improve brain functions in the PSD model.

## Introduction

Ischemic stroke is a sudden loss of brain blood supply, which can cause long-term neuropsychiatric symptoms, including depression, anxiety, mania, and associated cognitive disorders ^1^. Developing medications to support post-stroke brain recovery is one of the main priorities for reducing the economic burden on families and society. Post-stroke depression (PSD) is a major neuropsychiatric disorder induced by the stroke insult and seriously affects the functional recovery of the brain ^2^. Maintaining psychiatric health can also promote brain repair. It has been reported that antidepressant treatment of stroke patients improves brain functional recovery ^3^. One of the underlying neurobiological factors is adult neurogenesis. In the hippocampus, adult neurogenesis is a key element of structural plasticity that plays a crucial role in regulating antidepressant behaviors ^4^. Neural stem cells (NSCs) continuously generate neural progenitors and develop into mature functional neurons. Declining neurogenesis leads to a suppressed ability to cope with psychiatric stress and subsequently results in the development of depression ^5^. In rodent depression models, hippocampal neural stem/progenitor cells (NSPCs) display significantly suppressed growth and differentiation ^6,7^. Promoting adult hippocampal neurogenesis can help improve cognitive function, as demonstrated by enhanced performance in Y-maze tests. Recent breakthroughs in single-cell transcriptomics now enable detailed examination of cell-type-specific alterations in neuropsychiatric disorders. Research on depression demonstrates that its single-cell profiles exhibit the most pronounced genetic dysregulation in deep-layer excitatory neurons and immature oligodendrocyte precursor cells^8^. Thus, neurogenesis could be a promising therapeutic target for preventing PSD.

The proliferation and differentiation of neural stem cells (NSCs) in the hippocampus are regulated by multiple factors ^9^. Identifying effective targets for regulating neurogenesis could promote drug development for functional brain recovery. Metabolic factors act as key regulators of adult neurogenesis ^5^. PI3K/GSK3 signaling plays multiple roles in regulating neural plasticity, including synaptic plasticity, neurogenesis, and synaptogenesis ^10,11^. PI3K is an upstream signalling kinase that inhibits GSK3, which is a downstream kinase with diverse cellular effects with multiple cellular effects ^12^. Wnt signaling, a key regulator for adult neurogenesis, functions as the therapeutic pathway to improve post-stroke brain functions via the GSK3 pathway ^13^. Given the significant influence of metabolic pathways on adult neurogenesis regulation, metabolic compounds show promise as neuroprotective agents. These agents could mitigate neural damage and enhance regeneration processes. Methyl gallate (MET), a naturally occurring phenolic ester formed by gallic acid methylation, demonstrates this potential. Found abundantly in numerous plant species and traditional Chinese medicinal botanicals, this compound exhibits broad-spectrum bioactive properties relevant to neural protection - a capacity particularly evident in ischemic stroke, where MET functions as a well-established Chinese herbal compound with neuroprotective effects. ^14,15^. Previous reports indicate that MET can protect the brain against methamphetamine-induced neurodegeneration by mediation of GSK3 signaling pathways ^16^. Moreover, chronic MET treatment improves post-stroke angiogenesis and recovery after experimental stroke ^14^. One of the mechanisms of action of MET is the activation of AMPK ^17^, which also plays an important role in regulating animal behavior and adult neurogenesis ^18^. However, little is known about whether MET treatment could attenuate PSD-induced brain dysfunctions and if such effects are associated with activating AMPK, PI3K, or GSK3 kinases. To address such questions, computer-simulated molecular docking technology offers an efficient approach for investigating drug-target interactions, as demonstrated by Zhou Yang and colleagues, who used this method to uncover crosstalk between echinacoside and Nrf2, thereby proposing a molecular mechanism for echinacoside’s improvement of PSD ^19^.

In this study, we identified unique cellular profiles and molecular characteristics of PSD based on single-cell transcriptomics analysis and explored the association between MET and AMPK/GSK3 through molecular docking. Subsequently, we established a post-stroke depression mouse model and investigated the effects of MET in behavioral protection, neurogenic promotion, and the underlying cell signaling mechanisms.

## Materials and Methods

### Animals

C57BL/6N male mice were obtained from the Laboratory Animal Unit of Shanxi Medical University. All animals were kept with standard temperature (25±2°C) and humidity (60±5%) with ad libitum feeding. All animal experimental procedures were approved by Shanxi Medical University (IACUC2018-17). Anesthesia was administered using nose cone-delivered isoflurane (maintained at 1.5% in 80% N_2_O and 20% O_2_). A silicon-induced MCAO rubber-coated 7-0 monofilament (Doccol Corporation, Redlands, CA) was placed in the internal carotid artery, after which the monofilament was advanced to occlude the MCA. The filament was withdrawn 45 min after occlusion, and reperfusion was confirmed by laser Doppler monitoring. The surgical wound was sutured, and the mice could recover from anesthesia. Mice were anesthetized with halothane (1.5 to 2% in O2-enriched air by face mask) and kept warm on water pads.

After a 5-day habituation, mice were then subjected to mild stress for 2 weeks by restraint for 6 hours per day to induce post-stroke depression. Treatment groups received MET (low dose: 35 mg/kg, high dose: 100 mg/kg) daily by oral gavage. MET treatment alone was performed at 100mg/kg/d for 2 weeks. Bromodeoxyuridine (BrdU, 50 mg/kg/day) was administered to mice with i.p. (Intraperitoneal) injection (Fig. 1). For LY294002 inhibitor treatment, LY 294002 (4 mg/kg/d, 1 mg/ml in saline) and compound C (10mg/kg/d, 2.5 mg/ml in saline) were used for treatment along with 100 mg/kg i.p. MET-treatment group.

**Fig. 1:**
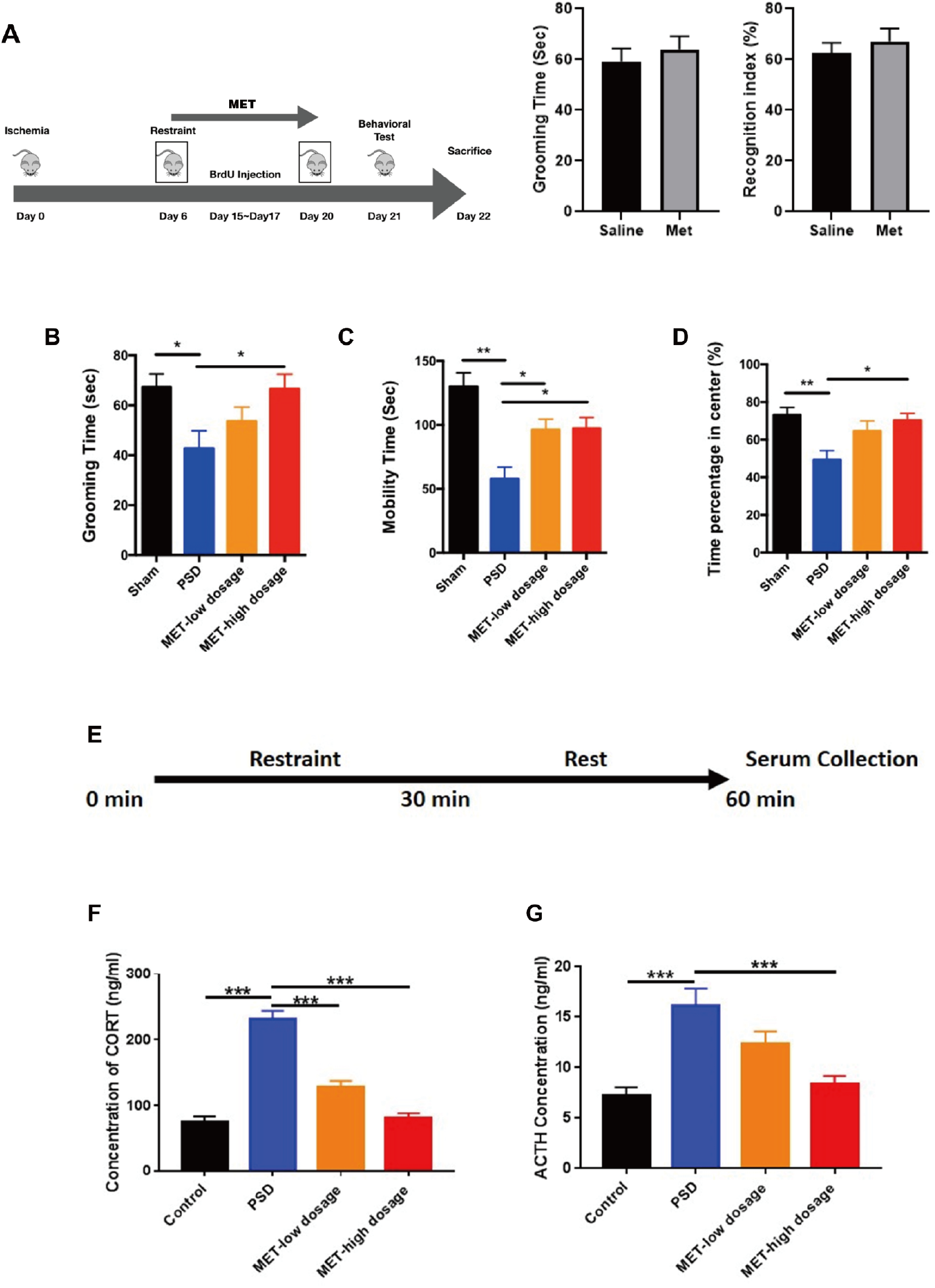
MET attenuated PSD-induced depressive-/anxiety-like symptoms. **(A)** Experimental procedure of PSD establishment and MET treatment. n=8 per group. **(B)** Grooming time in the plash test. (**C)** Overall mobility time in FST. (**D)** Time spent in the central region of the field in OFT. (**E)** Experimental procedure to perform mild stress and collect a serum sample. (**F)** Serum levels of CORT. (**G)** Serum levels of ACTH.

### Behavioral tests

#### Splash test

The splash test was performed according on a previous report ^20^. This test was conducted by spraying a 10% sucrose solution on the back of mice in their home cage. The sucrose solution dirties the coat and induces grooming behavior due to its viscosity.

#### Forced swim test

Mice were placed in a cylinder water tank (30 cm height × 20 cm diameter) with the water temperature 23~25°C for 6min. The whole behavior test process was video-recorded, and mobility time within the last 4 minute (define the first 2 minute as habituation), including struggling and free swim time, was recorded for analysis.

#### Open field test

Mice were put into an open field with a size of 43cm × 43cm for 10min. Mice could freely move in the arena, which was artificially separated by 16 equal squares. The duration of the animal’s travel in the central area was recorded to analyze the anxiety level. The open field was cleaned with 75% ethanol before the test. Mice used for emotional behavioral tests, including the splash, forced swim test, and open field test, were a different group from the mice used in cognitive tests, including the water maze and objective recognition task.

#### Morris water maze

Morris water maze (MWM) test began with an acquisition task to assess the spatial learning behavior. Briefly, each mouse was placed in the water for 60 seconds to freely search for the hidden platform and form spatial memory in reference to the marks on the walls of the water tank. The escape latency was recorded, and the acquisition task was performed for 4 consecutive days. Probe trail was acted in the day after the acquisition task by removing the platform. Each mouse was then put into the water for a 60-seconds free swim. Swimming time in the original platform zone was recorded to evaluate the learning ability. Visible platform test was conducted to assess any impairment in visual or motor ability induced by treatments.

#### Objective recognition task

After the MWM task, mice performed an object recognition task (ORT). Mice were first habituated to the ORT for 10 min, during which the total distance travelled was recorded to evaluate the interference with motor ability. During familiarization, mice were exposed to an open field with two identical objects (two red cubes) for 10 minute or stopped when the total exploration time reached 20 seconds. After a two-hour interval, mice were returned to the open field with one familiar object (red cube) and a novel object (blue cylinder) for 10 minute or until the total exploration time reached 20 seconds. The object recognition index was recorded with (novel exploring time/total exploring time) and calculated as a percentage.

### ELISA

Serum corticosterone and ACTH (adrenocorticotropic hormone) concentrations were detected by ELISA kit (Abcam, UK; Enzo, US). Animals in all groups after behavioral tests was subjected to restraint stress for 30 minutes, and blood was collected 30-minute after the end of stress (Fig. 4A). Serum was prepared by centrifuging the blood sample at 1500×g for 15 min. Experimental procedures followed previous reports ^21,22^. Concentrations and standard curves were determined by a plate reader (Bio-Rad) at a wavelength of 450 nm.

### Cell Culture

Primary NSCs culture and differentiation were conducted following previously described ^23^. Briefly, fetal Sprague–Dawley rats (embryonic days E14-15). The dissociated cells (1 × 105 cells/ml) were suspended in DMEM/F12 medium replenished with 2% B27 supplement, recombinant human bFGF (20 ng/ml), and EGF (20 ng/ml) with B27 applied. For the NSCs differentiation study, 2–5 passages of NSCs were mechanically dissociated as single cells and directly plated onto poly-L-lysine coated coverslips in the culture medium, withdrawn growth factors for 7 days, and started the 20μM MET and LY294002 (35 μM) or vehicle, along with a hypoxia protocol. Briefly, cells were placed into a chamber flushed with 2.5% O_2_ plus 5% CO_2_ balanced with 92.5% N_2_ and maintained in the hypoxia chamber at 37 °C for 4 days. Cells after treatment were fixed with 4% PFA.

### Immunofluorescence

Mice were anesthetized by Ketamine (200 mg/kg) and Xylazine (10mg/kg), and sacrificed by cardiac perfusion with 4% paraformaldehyde (PFA). Hippocampal cryosections and cell cultural slices were prepared on cover slips. Primary antibodies (Mouse-anti-BrdU, CST, 1:400; Rabbit-anti-DCX, CST, 1:400; Rabbit anti-Ki67, CST, 1:400; Mouse anti-Nestin, Abcam, 1:200; Mouse anti-Sox2, Abcam, 1:300) were incubated with sections at 4°C overnight. Secondary antibodies (Goat-anti-Rabbit-Alexa fluor 488, ThermoFisher, 1:800; Goat-anti-Mouse-Alexa fluor 568, ThermoFisher, 1:800) were incubated with sections for 2 h at RT. DAPI (Sigma) was incubated with sections for 10min. Tissue IF images were obtained with confocal microscopy (Nikon, C2^+^) with Z-stack for 20μm and projection on Z-stack was performed with ImageJ.

DAPI (Sigma) was incubated with sections for 10min. Tissue IF images were obtained with confocal microscopy (Nikon, C2^+^) with Z-stack for 20μm and projection on Z-stack was performed with ImageJ.

### Western blot

Protein lysate of cultural cells after treatment was prepared. Equal protein amounts were resolved by SDS-PAGE and transferred to polyvinylidene difluoride (PVDF) membranes (Millipore, USA). Membranes were blocked for 1 hour at room temperature with constant shaking in Tris-buffered saline containing 0.1% Tween-20 (TBS-T) and 5% BSA. Subsequently, membranes were incubated overnight at 4°C with the following primary antibodies from Cell Signaling Technology (CST): Rabbit anti-pGSK3β (Y216) (1:1000), Rabbit anti-GSK3β (1:1000), Rabbit anti-pAMPKα (Thr172) (1:1000), and Rabbit anti-AMPKα (1:1000). After washing, membranes were incubated with Goat anti-Rabbit HRP-conjugated secondary antibody (1:5000) for 2 hours at room temperature. Results were analyzed using Quantity One (Bio-Rad, USA).

### Single-Cell Transcriptomic Analysis of PSD

We performed an in-depth analysis on a public available single-cell RNA sequencing dataset (GSE232936) from the hippocampal region of rats with PSD. The original dataset includes four experimental groups: NC (negative control, SRR24661533), MCAO (middle cerebral artery occlusion model, SRR24661532), DEPR (chronic stress-induced depression model, SRR24661531), and MD (post-stroke depression model, SRR24661530). Raw FASTQ files were aligned to the reference genome and quantified using Cellranger v9.0.1, and downstream single-cell analysis was conducted with Seurat v4.4.0. UMAP was used for dimensionality reduction and clustering, and cell clusters were annotated based on canonical marker gene expression. Subsequent analyses comprised three components: (1) Comparison of cell type proportions across experimental groups; (2) Differential expression analysis of GSK3/AMPK gene family members; (3) AUCell-based evaluation of neurogenesis/synaptogenesis pathway activities, with group-wise comparisons of activity scores. All statistical comparisons employed Wilcoxon rank-sum testing with significance thresholds defined as *P < 0.05, **P < 0.01, ***P < 0.001, and ****P < 0.0001..

### Molecular Docking Analysis of MET

We retrieved MET’s chemical structure from PubChem (CID: 7428) and acquired protein sequences for AMPK subunits (PRKAA1/2, PRKAB1/2, PRKAG1-3) and GSK3 isoforms (GSK3A/B) via UniProt. All available human and mouse structural conformations—including experimentally resolved PDB entries and AlphaFold DB v4 predictions—were incorporated. Using AutoDock, we conducted rigid-receptor molecular docking against these targets. Each protein-ligand pairing underwent 10 independent docking runs, with the conformation having the lowest binding energy (in kcal/mol) selected as optimal. Binding energies were categorized as follows: >-4 kcal/mol (weak), −4 to −7 kcal/mol (moderate), and <-7 kcal/mol (strong). PyMOL facilitated the visualization of the final binding pose.

### Statistics

For comparisons across multiple groups, statistical analyses were performed using a one-way analysis of variance (ANOVA) followed by a multiple comparison test using GraphPad Prism 5 (GraphPad Software Inc., CA, USA). For ADP and ATP analyses, a two-way ANOVA, following a multiple comparison test, was performed. A Student’s t-test was performed for two independent group comparisons. A p-value < 0.05 was considered statistically significant. All values are expressed as mean ± SEM.

## Results

### Methyl gallate attenuates depression-/anxiety-symptoms in the PSD model

We performed behavioral tests to evaluate the antidepressant effects of MET in the PSD mouse model (Fig. 1A). Additionally, an antidepressant assessment using the splash test and a cognitive behavioral assessment using ORT showed that MET treatment alone did not alter the behavioral performance (Fig. 1A). Compared with the PSD model, high dose (100 mg/kg) of MET treatment increased the grooming time in the splash test and prolonged mobility time in FST (Fig. 1B, C; one-way ANOVA, p<0.05 in both tests), indicating that MET attenuated depressive symptoms. Anxiety-like behaviors were assessed by performing OFT. The PSD group showed a significant reduction in time in the center area of the open field (Fig. 1D; p<0.01). Compared with the PSD group, the MET high-dose group showed an increased time in the center region of the open field (Fig. 1D; p<0.05). Collectively, the behavioral tests indicated that MET alleviated the depressive and anxiety-like behaviors in PSD mice.

We next assessed the activity of the hypothalamus-pituitary-adrenal (HPA) axis by measuring stress-induced production of corticosterone (CORT) and ACTH (Fig. 1E). After the 30-minute restraint and 30-minute recovery, concentrations of CORT and ACTH remained significantly higher in PSD mice compared with controls (Fig. 1F, G; p<0.001). Both doses of MET treatment markedly reduced the concentration of CORT and ACTH (Fig. 1F, G; p<0.001). The low dose of MET effectively reduced the serum level of CORT (Fig. 1F; p<0.001), but did not show a significant effect on serum ACTH (Fig. 1G; p=0.157). Overall, MET treatment could attenuate depression- and anxiety-like symptoms in the PSD model in a dose-dependent manner.

### Methyl gallate treatment attenuates the cognitive dysfunctions in the PSD model

We next conducted the cognitive tests to evaluate the effects of MET on cognitive function. The acquisition task in MWM was first performed to test spatial learning. Both doses of MET treatment significantly reduced the escape latency with the increasing training days (Fig. 2A, p<0.001 vs. PSD). In the probe trial, the swim time of mice in the target zone was also markedly prolonged by MET treatment, indicating that MET improved spatial memory (Fig. 2B, p<0.001 vs. PSD). The ORT result showed that MET treatment at the high dose significantly increased the recognition index in comparison with the PSD group (Fig. 2E, p<0.01). There were no significance difference in escape latencies in the visible platform test as well as the total distance moved during ORT habituation among groups, indicating the motor ability was not affected by the experiments (Fig. 2C, D). Collectively, these findings suggest that MET can also improve cognitive function in the PSD model.

**Fig. 2:**
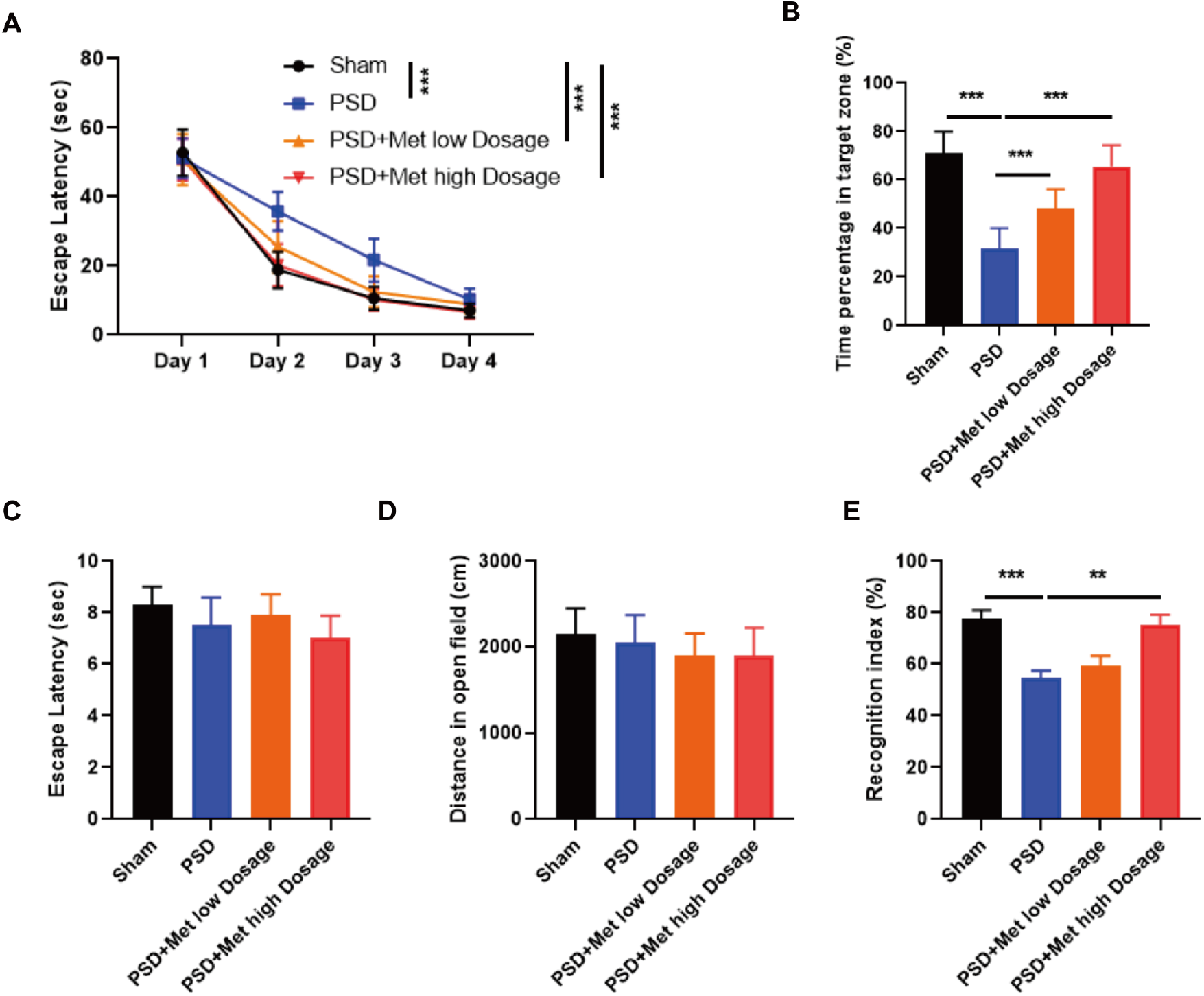
MET treatment improved cognitive performance in the PSD model. **(A)** Statistical analysis to show the effects of MET on escape latency on different training days. **(B)** Statistical analysis to show the difference in the percentage of time in the target zone in the probe trial of MWM. (**C, D)** Statistical analysis to show the escape latency in the visible platform and move distance in ORT habituation. **(E)** Statistical analysis to show the difference of the recognition index in ORT.

### The cellular types of PSD resemble those of depression but exhibit a more pronounced impairment of neurogenesis

We performed UMAP dimensionality reduction on 31,668 cells based on sample identity (Fig. 3A) and cell type classification (Fig. 3B). The results revealed that Microglia and Oligodendrocytes exhibited distinct separation, which is likely attributable to their unique transcriptional profiles (Fig. 3C). In addition to Microglia, several peripheral immune cell types (T cells, B cells, Macrophages, and Neutrophils) were detected, suggesting that under pathological conditions, these immune cells may infiltrate the brain parenchyma through the blood-brain barrier (BBB) and participate in neuroinflammatory surveillance and modulation. A comparison of cell-type composition across groups (Fig. 3D) demonstrated that the cellular distribution in the MD group closely resembled that of the DEPR group. Specifically, Microglia, Neutrophils, Plasma cells, and Astrocytes were more abundant in MD than in MCAO, indicating that the hippocampus in PSD is characterized by heightened inflammatory cell infiltration relative to MCAO.

**Fig. 3.**
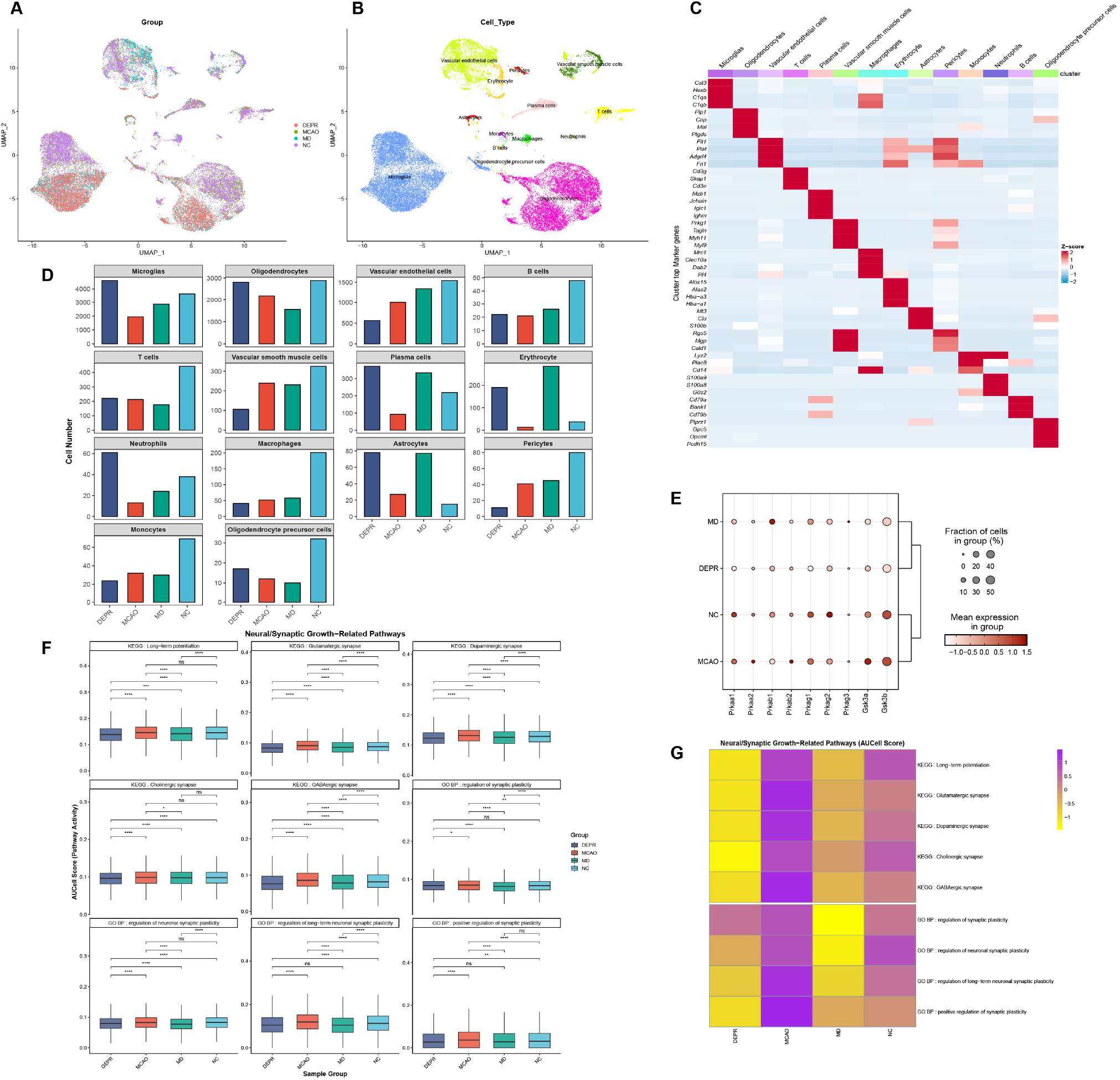
Single-cell transcriptomic analysis of the hippocampus in PSD rat models. **(A)** UMAP plot showing the distribution of cells from the four experimental groups. **(B)** UMAP plot illustrating cell type annotations. (**C)** Heatmap displaying the average expression levels of canonical marker genes used for cell type identification. (**D)** Comparison of cell counts across 14 identified cell types among the four groups. (**E)** Average expression levels of AMPK and GSK3 gene family members across groups. (**F)** Statistical comparison of AUCell scores for nine neurogenesis/synaptogenesis-related KEGG/GO signaling pathways among the four groups. Pairwise comparisons were performed using the Wilcoxon rank-sum test. Significance levels: P < 0.05 (*), P < 0.01 (**), P < 0.001 (***), P < 0.0001 (****). (**G)** Heatmap showing the average pathway activity scores of the nine neurogenesis / synaptogenesis-related pathways across the four groups.

**Fig. 4:**
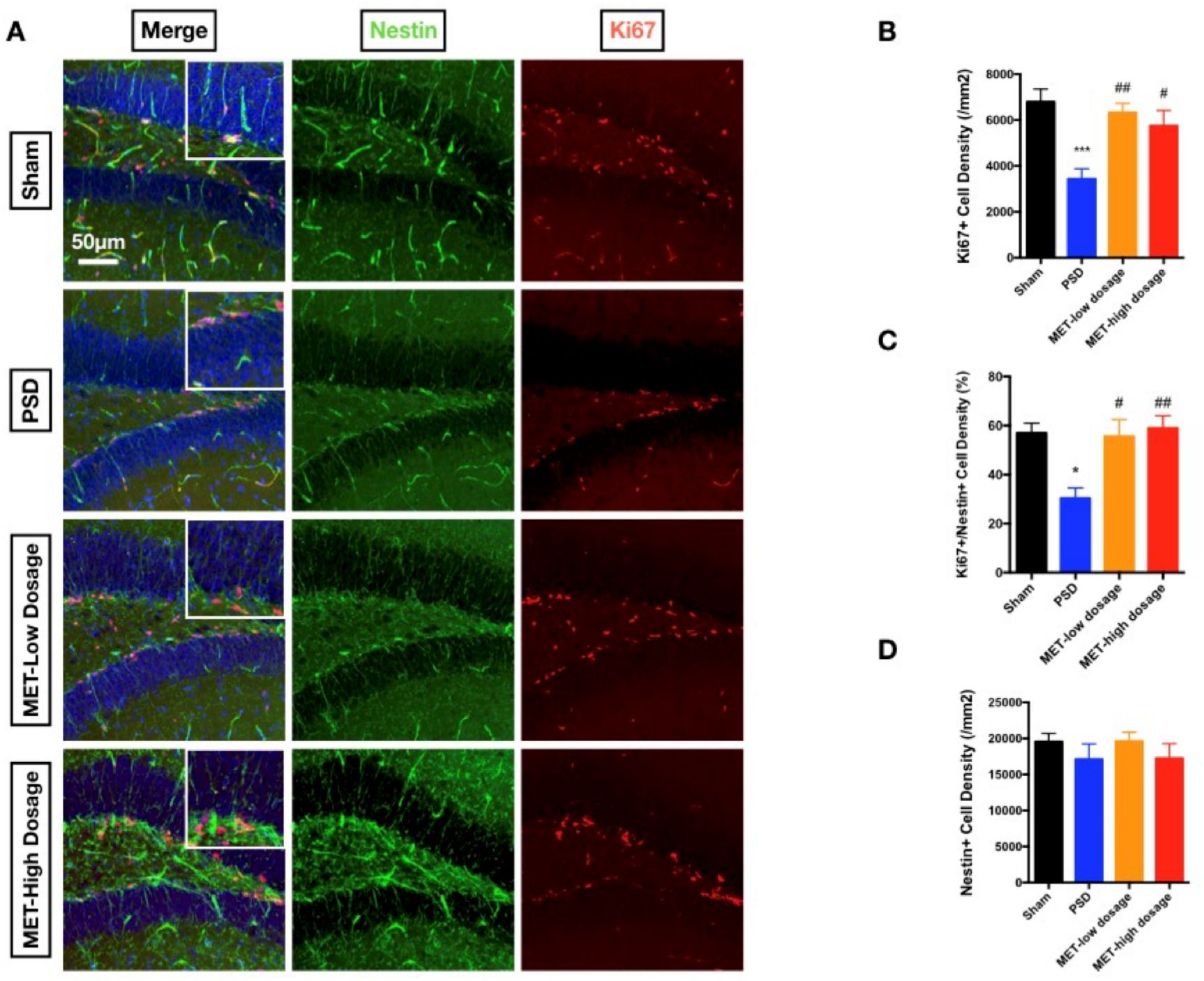
MET improved the proliferation of NSCs in the hippocampus. **(A)** IF image to present the density of newborn cells labelled with Nestin (Green) and Ki67 (Red). Nuclei were marked by DAPI staining (Blue). **(B)** Statistical analysis to show the Ki67^+^ cell density in different groups. (**C, D)** Statistical analysis to show the Ki67^+^/Nestin^+^ as well as Nestin^+^ cell density in different groups.

We then examined the expression patterns of the AMPK and GSK3 gene families across the four groups (Fig. 3E). Except for Prkab1, most genes exhibited similar expression levels in MD and DEPR yet differed from the expression profiles observed in MCAO and NC. For instance, Prkaa1 and Gsk3b were predominantly expressed in the NC and MCAO groups but showed markedly reduced expression in the MD and DEPR groups.

Furthermore, we evaluated the activity of signalling pathways related to neurogenesis and synaptic growth across the four groups. Boxplot analysis (Fig. 3F) revealed a significant reduction in the activity of most neural/synaptic pathways in the DEPR and MD groups compared to MCAO. Heatmap (Fig. 3G) analysis further confirmed that neurogenesis-related pathway activities were broadly suppressed in both DEPR and MD, with PSD exhibiting even lower activity in synaptic plasticity regulation.

These findings suggest that the pathological features of PSD are comparable to those of depression, characterized by enhanced hippocampal inflammation and impaired neurodegeneration. However, PSD demonstrates a further exacerbation in the suppression of synaptic plasticity, indicating more severe dysfunction in neural remodelling capacity.

### Methyl gallate treatment promotes adult hippocampal neurogenesis in the PSD model

We first assessed NSC proliferation by labeling Ki67^+^ and Ki67^+^/Nestin^+^ cell numbers. Compared with the PSD model, MET treatment strikingly restored the proliferation of NSCs marked by Ki67^+^/Nestin^+^ staining (Fig. 4, A-C). Total NSCs density marked by Nestin did not differ among groups (Fig. 4D), indicating the NSCs population was not affected in PSD. We then tested the generation of immature neurons marked with DCX. BrdU was injected 5 days before sacrifice. As a result, PSD decreased the density of BrdU^+^ cells in the hippocampus (Fig. 5A, B; p<0.01 compared to control). While MET treatment at both low and high doses significantly increased BrdU^+^ cell density compared with the PSD group (Fig. 5, A, B; p<0.05 for low dosage, p<0.01 for high dosage compared with the PSD group), indicating that MET enhanced the generation of NSCs in the hippocampus. BrdU^+^/DCX^+^ cells were analyzed to evaluate the newborn immature neurons. We observed the decreased BrdU^+^/DCX^+^ cells in the PSD group compared with the control (Fig. 5C; p<0.01 compared with the control). MET treatment with both dosages markedly improved the generation of immature neurons, as reflected by the increased density of dual-positive cells compared with the PSD group (Fig. 5C; p<0.05 for both dosages compared with the PSD group). Collectively, MET could improve the proliferation and neuronal differentiation of NSCs in the PSD model.

**Fig. 5:**
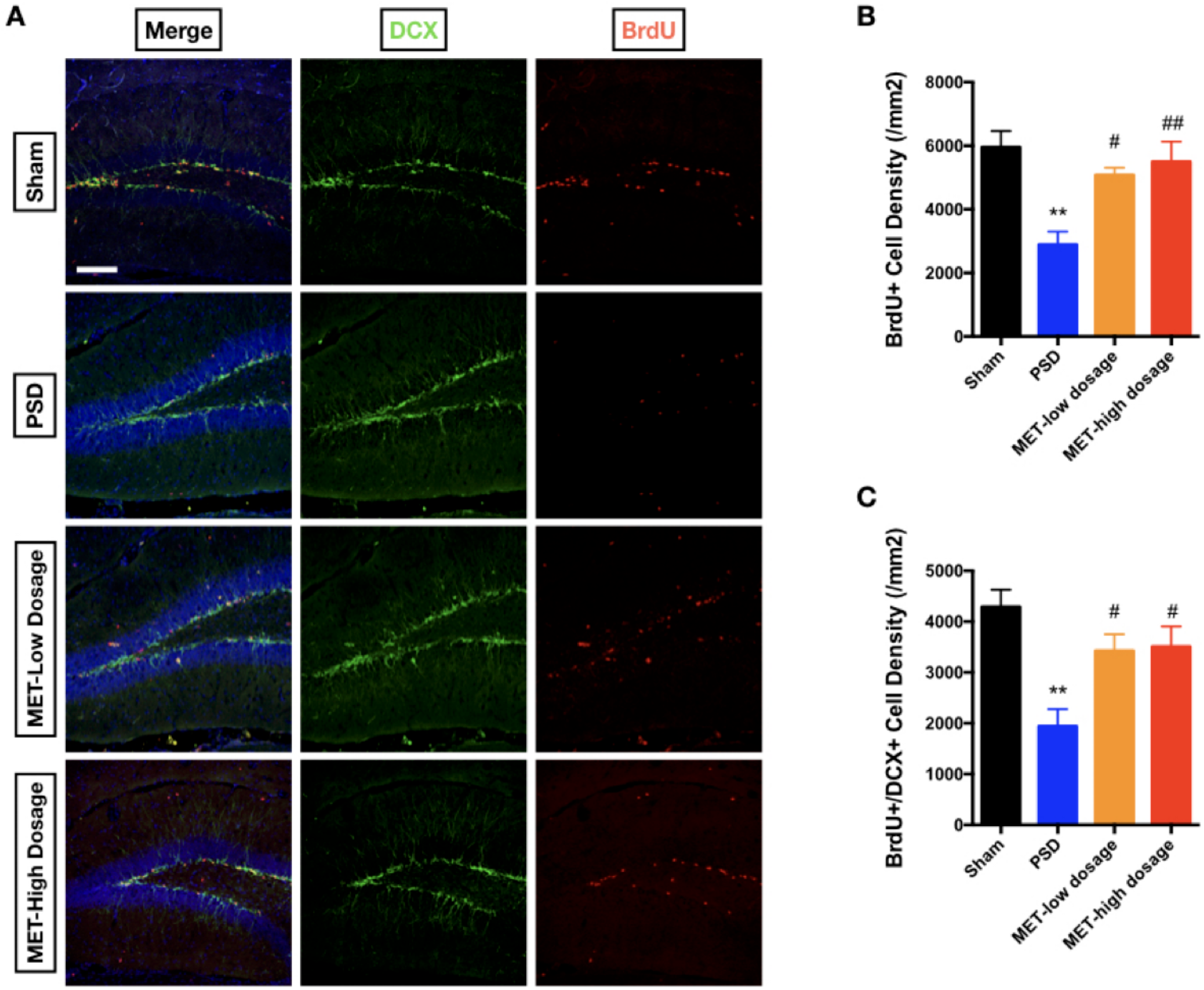
MET improved neuronal differentiation of NSCs in hippocampus. (**A**) IF image to present the density of newborn cells labelled with BrdU (Green) and neuroblasts (NB) by DCX (Red). Nuclei were marked by DAPI staining (Blue). **(B)** Statistical analysis to show the BrdU^+^ cell density in different groups. (**C)** Statistical analysis to show the cell density of dual-positive BrdU and DCX cells. Bar=100μm, n=5 per group.

### GSK3β/AMPK-mediated neurogenesis underlies the efficacy of MET in PSD

Molecular docking analysis revealed that MET exhibits potential interactions with most subunits of AMPK and GSK3, except for GSK3A (Supplementary Table 1). We visualized the binding pockets corresponding to the top-ranking docking results based on binding affinities. Among all subunits examined, MET exhibited its strongest interaction with GSK3B, displaying a binding energy of −5.81 kcal/mol (Fig. 6A). Subsequent analysis revealed progressively weaker binding affinities with: PRKAG1 (−5.53 kcal/mol, Fig. 6F); PRKAG3 (−5.53 kcal/mol, Fig. 6H); PRKAB2 (−5.22 kcal/mol, Fig. 6E); PRKAA2 (−4.84 kcal/mol, Fig. 6C); PRKAG2 (−4.82 kcal/mol, Fig. 6G); PRKAB1 (−4.77 kcal/mol, Fig. 6D); and PRKAA1 (−4.58 kcal/mol, Fig. 6B). These findings suggest that MET may interact with key components of the AMPK/GSK3 pathway and potentially mediate its pro-neurodegenerative effects.

**Fig. 6:**
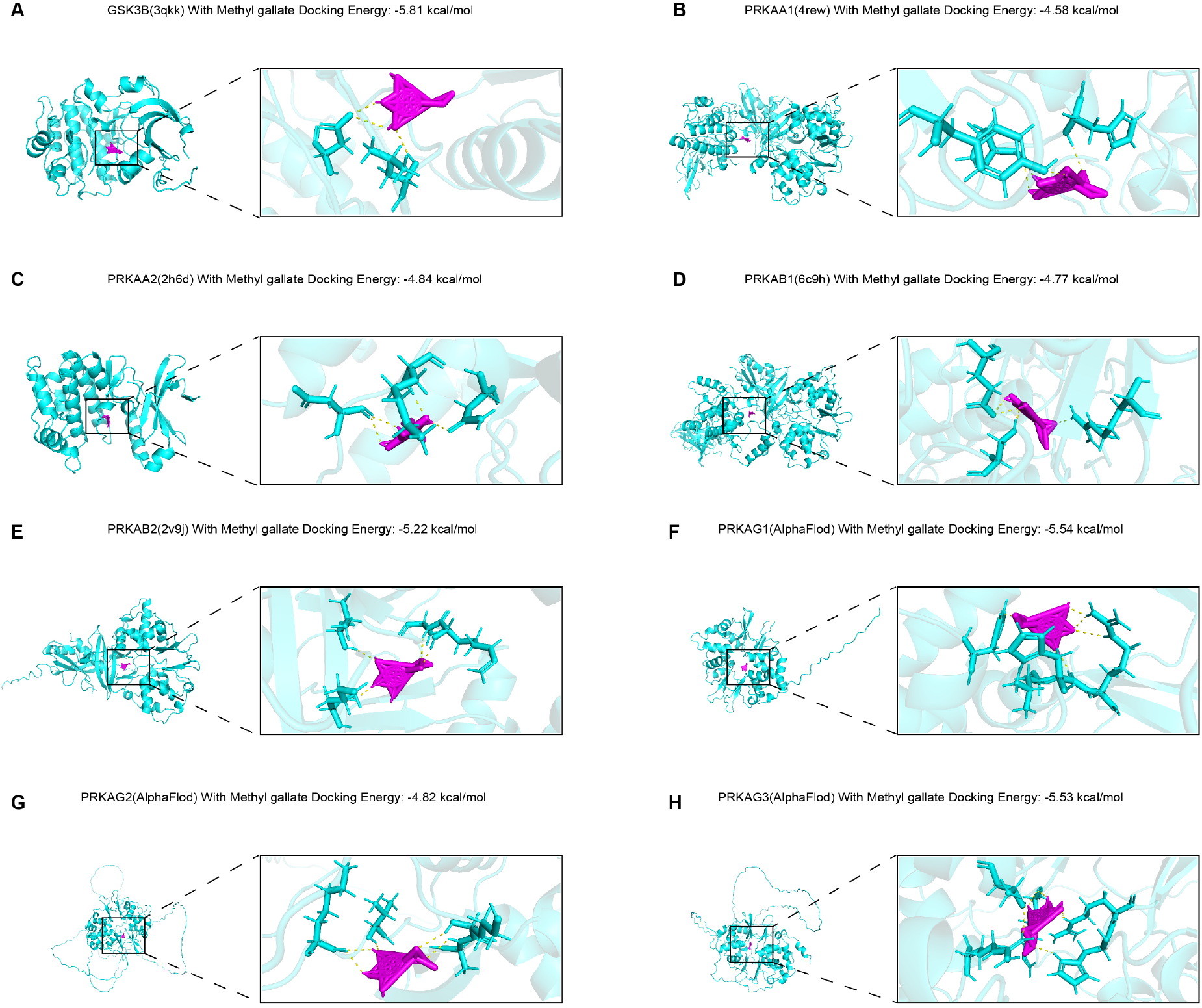
Molecular docking results of MET with AMPK/GSK3 gene family members. Among the GSK3 family, only GSK3B (**A)** showed binding affinity with MET. AMPK family members included PRKAA1 (**B)**, PRKAA2 (**C)**, PRKAB1 (**D)**, PRKAB2 (**E)**, PRKAG1 (**F)**, PRKAG2 (**G)**, and PRKAG3 (**H)**.

In hippocampal tissue, we found that MET treatment dramatically decreased the phosphorylation of GSK3β at Y216 and increased the phosphorylation of AMPK at Thr172 (Fig. 7B, C; p < 0.001, MET vs. PSD group). Herein, we assumed that the activation of the PI3K-Akt pathway and the activation of AMPK might be the key mechanisms underlying the pro-neurogenic effects of MET. To confirm this hypothesis, cultural neural progenitors were exposed to hypoxia (Fig. 7A). Under hypoxia, the effect of MET to enhance DCX+ neural progenitor differentiation was compromised by the PI3K inhibitor LY294002 and the AMPK inhibitor compound C, separately, which indicates the critical role of GSK3 and AMPK in MET-mediated neural development. To further confirm the roles of GSK3 and AMPK in MET-mediated behavioral effects, we treated the PSD model with LY294002 and compound C, along with MET injection (Fig. 7E). Both treatments significantly inhibited the effects of the MET to prolong the grooming time in the splash test and to restore the recognition index in ORT (Fig. 7F, G; p<0.01 in splash test, p<0.001 in ORT). The combination of the two inhibitors’ administration also presented an inhibitory effect on the performance of mice in the splash test and ORT. However, the combination treatment showed no significant difference compared to either Compound C or LY294002 (Fig. 7F, G). This result suggests that the activation of PI3k and the resulting inhibition of GSK3β, along with the activation of AMPK, are important in mediating the effects of MET on pro-neurogenic and behavioral improvement.

**Fig. 7:**
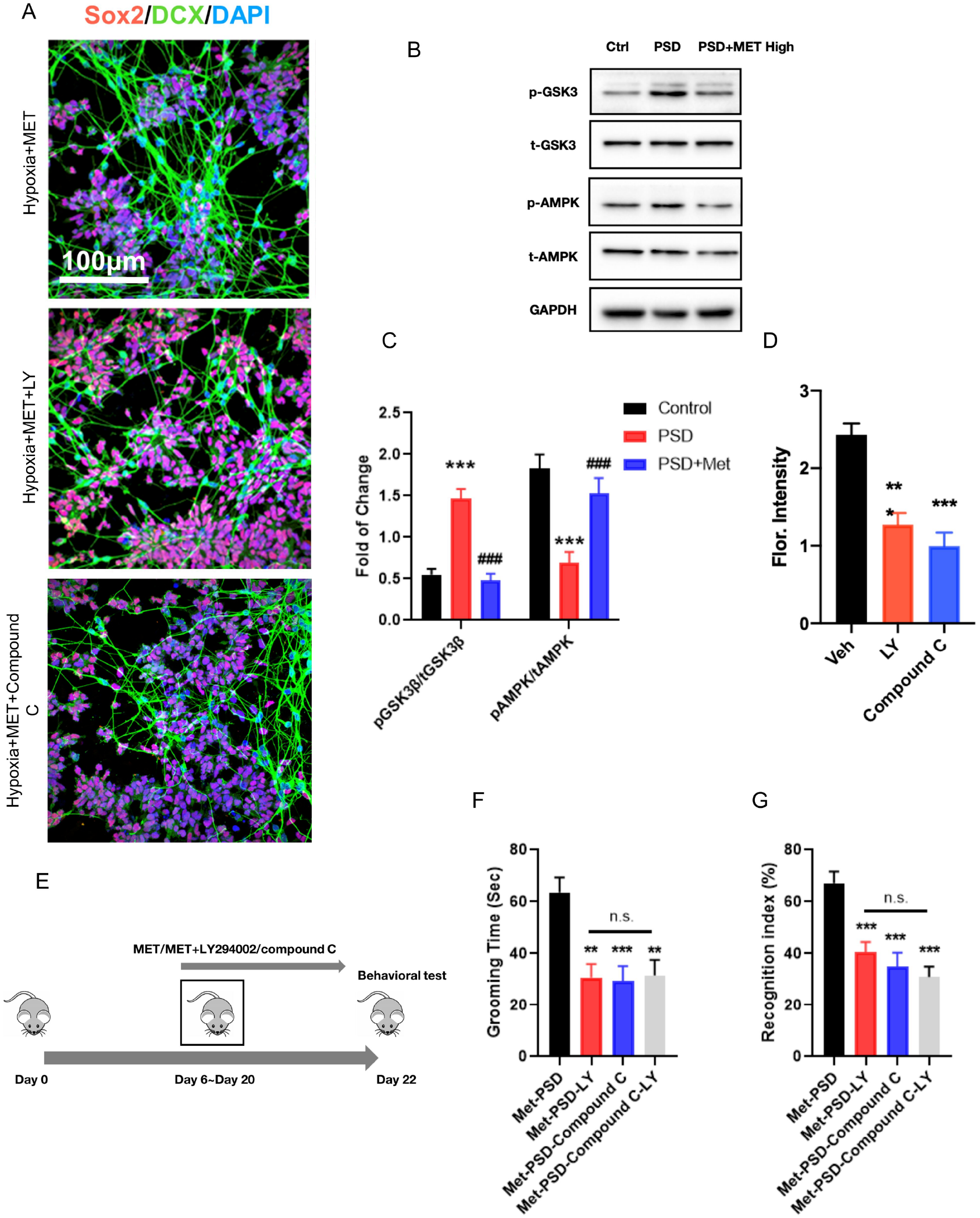
Neurogenesis is increased by MET in the hippocampus. **(A)** Levels of NSCs marker Sox2 (Red) and immature neural fiber marker DCX (Red). (**B, C)** Western blot bands and statistical analysis to show the phosphorylation of GSK3β on Y216 and the total GSK3β protein as well as the phosphorylation of AMPK on Thr172 and total protein level in hippocampal tissue. (**D)** DCX levels in cultural neural induction experiments. (**E)** Experimental procedure to show the LY294002 or compound (C) treatment in the MET group. (**F, G)** Grooming time in the splash test and recognition index in ORT.

## Discussion

Attenuating post-stroke depression (PSD) is critical for promoting functional brain recovery after a stroke. A recent study profiled hippocampal endothelial cells, microglia, and oligodendrocytes in a PSD model, showing evidence of distinct alterations and enhanced cellular interactions unique to PSD ^24^. Through in-depth single-cell analysis, we observed that PSD shares cellular and molecular similarities with primary depression. Notably, PSD exhibits a more pronounced downregulation of key regulators (e.g., PRKAA1 and GSK3β), resulting in more significant suppression of neurogenesis and synaptic plasticity-related pathways. This finding indicates severely compromised hippocampal neural repair capacity in PSD, providing a mechanistic basis for the development of neurodegenerative therapies. Here, we report a previously uncharacterized role of MET in attenuating PSD-associated emotional and cognitive dysfunction by enhancing hippocampal neurogenesis. GSK3β inhibition by MET represents the primary mechanistic basis for this therapeutic effect.

In previous reports, the routine period for depression models is usually not less than 1 month^25^. In the present study, ischemic stroke reduced the adaptability of mice to psychiatric stressors. Following ischemia, 12 days of restraint stress induced depressive symptoms and a hyperactive HPA axis (Fig. 1). Previous reports demonstrated that MET is effective in attenuating depression-associated symptoms in a stroke animal model ^14^. Daily treatment with 100 mg/kg MET significantly improved the antidepressant behaviors in the PSD model. These data expand the application of MET as a neuroprotective agent. In the MWM test and the ORT, MET treatment significantly restored the learning and memory performance of the mice, indicating that MET could also improve cognitive functions. Most PSD animal models show reduced neurogenesis in the hippocampus ^26^. Promoting neurogenesis is widely considered a key strategy for preventing PSD and improving post-stroke recovery. In the present study, we observed that MET could enhance hippocampal neurogenesis by elevating the proliferation of NSCs and the production of newborn neurons in the hippocampus. Maintaining a minimum density of immature neurons is a prerequisite for the brain to clear anxiety-related memory and elevate memory resolution to prevent the development of anxiety ^27^. Previous results indicate that MET could promote synaptic plasticity in the high-fat diet (HFD) model ^28^, which further confirms the multiple regulatory functions of MET in the hippocampus. In addition to neurogenesis, synaptic plasticity also contributes to the regulation of antidepressant and cognitive behaviors ^29^. It could also be another consideration of the biological underpinning of MET in treating PSD behavioral deficits. Combined with the results from this study, MET might be a useful medication to improve hippocampal neural plasticity and thus benefit the post-stroke recovery.

It has previously demonstrated that corticosterone-induced depressive and anxiety-like behaviors can be attenuated by increasing neural regeneration alone in the hippocampus ^30^. Many antidepressants or cognitive-enhancing drugs, including fluoxetine, ketamine, and piracetam, have been reported to improve adult hippocampal neurogenesis ^31–33^. In the present study, we detected an increased stress response in the PSD model, reflected by elevated CORT and ACTH secretion. Increased CORT levels play a critical role in limiting the proliferation of hippocampal NSCs ^34^. Consistent with this, we also observed decreased activity of NSCs in the hippocampal DG region in the PSD model. MET treatment attenuated the stress response in parallel with enhancement of NSC proliferation, indicating that hippocampal neurogenesis is an important biological mechanism underlying the effects of MET in the PSD model. Moreover, we found that MET increased the number of newborn neurons, which likely derived from the increased proliferation of the NSCs. MET has also been shown to promote NSCs proliferation and neural differentiation in the hippocampal CA1 region of the acute ischemia/reperfusion model ^35^, supporting its therapeutic value of MET in stroke therapy.

Metabolic factors including insulin signaling, insulin-like growth factors (IGFs), adiponectin, and incretins, have been reported to regulate adult neurogenesis and related behaviors ^36–39^. Insulin activates the PI3K-Akt cell signaling pathway which controls cell growth and neurogenesis ^40^. Akt activity inhibits of GSK3β pathway that is linked to apoptosis. In the CNS, inhibiting GSK3 can help reduce neurodegeneration ^41–43^. GSK3 is a crucial neural developmental mediator that integrates multiple signaling pathways, including disrupted in schizophrenia 1 (DISC1), partitioning defective homolog 3 (PAR3), PAR6, and Wnt proteins^44^. It has been demonstrated that voluntary exercise increases adult hippocampal neurogenesis by increasing GSK3β activity in mice ^45^. It has also been reported that GSK-3β plays an important role in controlling autophagy induction by modulating the activation of the LKB1-AMPK pathway after serum deprivation, indicating the crosstalk between these two signaling agents ^46^.

Building on evidence that GSK-3β orchestrates LKB1-AMPK-dependent autophagy during metabolic stress, we posited that rectifying AMPK/GSK3 dysfunction could address neurodegenerative failure in PSD. While previous studies have employed molecular docking to explore microbial target interactions of plant-derived compounds such as MET and catechin 3-O-gallate ^47^, such structure-based strategies have rarely been applied to neurodegenerative targets. Here, we innovatively applied in silico docking to examine the direct binding potential between MET and core regulators AMPK and GSK3. Computational docking revealed high-affinity binding (binding energy ≤ −5.5 kcal/mol) between MET and conserved domains in AMPK and GSK3 regulators. This topological complementarity provides a mechanistic justification for MET-mediated AMPK/GSK3 modulation, supporting its pro-neurodegenerative efficacy observed in PSD models. Therefore, we will validate these predicted binding interactions through *in vitro/in vivo* assays, specifically probing MET-induced stabilization of AMPK/GSK3β complexes within hippocampal neurons under pathophysiological PSD-relevant conditions.

Here, we found that MET inhibited GSK3β in hippocampal tissue. By inhibiting PI3K, we achieved sustained activation of GSK3 (Fig. 7). This approach blocked the effects of MET on promoting neurogenesis and improving emotional and cognitive behaviors. Clinical trials have indicated that the GSK3 inhibitor lithium enhances motor recovery after stroke when administered early at a low dose ^48^. AMPK is a critical signaling kinase activated by Methyl gallate and thereby exerts multiple therapeutic effects ^49^. It is also possible that AMPK and GSK3 may be affected by MET in other brain regions, including the cortex and hypothalamus, which are also associated with the development of depression and post-stroke brain dysfunction ^50,51^. Normalizing serum glucose levels might additionally improve the AMPK activity in the hippocampus ^52^. This mechanism might also contribute to the effects of MET on PSD-induced behavioral deficits. In our results, we also found that administration of AMPK inhibitor compound C produces the same effects as LY294002 did (Fig. 7). These findings indicates AMPK activation as well as GSK3 inhibition are both necessary for MET to perform the pro-neurogenic and behavioral regulatory effects in the PSD model.

In conclusion, our study identified the new function of MET in preventing PSD-associated emotional and cognitive symptoms by promoting hippocampal neurogenesis. Inhibition of GSK3 as well as activation of AMPK with MET, could promote adult neurogenesis in hippocampus is the critical underlying mechanism. This study provides experimental evidence for the future application of metabolic drugs in brain functional recovery, not only in post-stroke conditions, but also in other neurodegenerations or psychiatric diseases. In future studies, direct MET regulation of GSK3β (vs. AMPK) requires validation with knockout models. Synaptic plasticity requires electrophysiological and protein data. Clinically, MET’s blood-brain barrier permeability and optimal human dosing, based on murine data, remain translational challenges.

Moreover, single-cell profiling revealed conserved cellular features between post-stroke depression (PSD) and primary depression, but PSD demonstrated more pronounced synaptic repair deficits and heightened neuroinflammation. Molecular docking confirmed the efficient binding of MET to AMPK/GSK3β targets (binding energy ≤ −5.5 kcal/mol), providing a mechanistic basis for its neurofunctional restoration in PSD.The mechanistic engagement of MET with neurodegenerative pathways establishes the compelling rationale for its deployment against PSD-associated neural damage. Our data demonstrate significant rescue of hippocampal neurogenesis deficits, though concurrent neuroinflammatory processes— particularly microglia-driven neurotoxicity and IL-6/TNF-α dysregulation—constitute unexplored moderators of therapeutic efficacy. Future work will employ spatial transcriptomics and fate-mapped microglia to resolve how MET-mediated repair mechanisms interface with neuroinflammation. Deciphering these interactions is expected to inform rationally designed combinational therapies that co-optimize neuroregeneration and inflammation resolution.

## Acknowledgment

This study was supported by Shanxi Scholarship Council of China (2016-051), Shanxi Natural Science Foundation (202103021224449), China National Natural Science Foundation (81300487), Yunnan Provincial Science and Technology Department (202402AA310033), Shenzhen Medical Academy of Research and Translation (D2401026), and Shenzhen-Hong Kong Cooperation Zone for Technology and Innovation (HZQB-KCZYB-2020056).

